# Cascading regime shifts within and across scales

**DOI:** 10.1101/364620

**Authors:** Juan C. Rocha, Garry Peterson, Örjan Bodin, Simon A. Levin

## Abstract

Regime shifts are large, abrupt and persistent critical transitions in the function and structure of systems (*1*, *2*). Yet it is largely unknown how these transitions will interact, whether the occurrence of one will increase the likelihood of another, or simply correlate at distant places. Here we explore two types of cascading effects: domino effects create one-way dependencies, while hidden feedbacks produce two-way interactions; and compare them with the control case of driver sharing which can induce correlations. Using 30 regime shifts described as networks, we show that 45% of the pair-wise combinations of regime shifts present at least one plausible structural interdependence. Driver sharing is more common in aquatic systems, while hidden feedbacks are more commonly found in terrestrial and Earth systems tipping points. The likelihood of cascading effects depends on cross-scale interactions, but differs for each cascading effect type. Regime shifts should not be studied in isolation: instead, methods and data collection should account for potential teleconnections.

## Introduction

Regime shifts occur across a wide range of social-ecological systems, they are hard to predict and difficult to reverse (*3*, *4*), while producing long term shifts in the availability of ecosystems services (*5*). A regime shift means that a system has moved from one stability domain to another, implying a shift from one set of dominating processes and structures to another set (*2*, *6*–*8*). Changes in a key variable (e.g. temperature in coral reefs) often makes a system more susceptible to shifting regimes when exposed to shock events (e.g. hurricanes) or the action of external drivers (*9*). Over 30 different regime shifts in social-ecological systems have been documented (*10*), and similar non-linear dynamics are seen across societies, finance, language, neurological diseases, or climate (*11*, *12*). As humans increase their pressure on the planet, regime shifts are likely to occur more often and severely (*13*).

A key challenge for science and practice is that regime shifts, in addition to being hard to predict, can potentially lead to subsequent regime shifts. A variety of cascading regime shifts have been reported (See Table S1). For example, eutrophication is often reported as a preceding regime shift to hypoxia or dead zones in coastal areas (*14*). Similarly, hypoxic events have been reported to affect the resilience of coral reefs to warming and other stressors in the tropics (*15*). If, why and how a regime shift somewhere in the world could affect the occurrence of another regime shift, however, remains largely an open question and a key frontier of research (*16*, *17*) that we here aim to address.

Research on regime shifts is often confined to well-defined branches of science, reflecting empirical, theoretical (*18*) or predictive approaches (*9*, *19*). These approaches require a deep knowledge of the causal structure of the of system or a high quality of spatio-temporal data. Hence research on regime shifts has generally focused on the analysis of individual types of regime shifts rather than potential interactions across systems. This paper takes another approach and instead explores potential interactions among a large set of regime shifts. We propose and investigate two types of interconnections: domino effects and hidden feedbacks. Domino effects occur when the feedback processes of one regime shifts affect the drivers of another, creating a one-way dependency (*9*, *17*, *20*). Hidden feedbacks rise when two regime shifts combined generate new (not previously identified) feedbacks (*16*, *20*); and if strong enough, they could amplify or dampen the coupled dynamics. Furthermore, we contrast these cascading effects, where the occurrence of a regime shift give rise to subsequent regime shifts, with the potentially multiplying albeit different effect of two regime shifts being caused by common drivers. Sharing drivers is likely to increase correlation in time or space among regime shifts, but not necessarily interdependence (*13*, *17*).

### Hypotheses of cascading effects

Domino effects occur when a variable that belongs to a feedback mechanism in a regime shift acts as driver in another. A feedback mechanism is a self-amplifying or dampening process characterized by a pathway of causal processes that return to its origin creating a cycle. Theory often treats drivers as slow variables, which assumes that their change are relatively slower than changes in state variables (*21*, *22*). Therefore, we expect that domino effects will be mostly dominated by connections from regime shifts that occur at larger spatial scales with slower temporal dynamics to regime shifts that operate at smaller spatial scales with faster temporal dynamics. Conversely, we expect that hidden feedbacks will occur when scales match (both in space and time), because for a new feedback to emerge they need to be somewhat aligned in the scale at which the process operate. Additionally, we expect that drivers sharing is more context-specific; in this case, regime shifts occurring in similar ecosystem types or land uses will be subject to relatively similar sets of drivers, increasing the likelihood of driver sharing.

Discovering how regime shifts might be interconnected requires a method for studying patterns in relational data. We analysed regime shifts as networks of drivers and feedback processes underlying their dynamics. These directed signed graphs allow us to explore driver co-occurrence, directional pathways and emergent feedback cycles of coupled regime shift networks (See SM Methods). The empirical basis for our investigation draws from the regime-shifts database(*10*), to our knowledge the largest online repository of regime shifts in social-ecological systems (Fig S2, SM Methods). It offers syntheses of over 30 types of regime shifts and > 300 case studies based on literature review of > 1000 scientific papers (*10*). The database consistently describes regime shifts in terms of their alternative regimes, drivers, feedback mechanisms, impacts on ecosystem services, and management options. It provides a set of 75 categorical variables (Table S2) about impacts, scales, and evidence types (*10*) that we used to test our hypotheses. Regime shifts are also described using consistently coded causal-loop diagrams as a summary of the drivers and underlying feedbacks of each regime shift (Fig S1). We converted the causal diagram for each regime shift into a network by creating the adjacency matrix *A*, where *A*_*i*,*j*_ is 1 if there is a connection or zero otherwise. A link between two nodes in these networks means that there is at least a scientific paper reviewed in the database providing some evidence for such causal relationship (*13*). Link sign *W*_*i*,*j*_ represents a link polarity taking –1 if the relationship is expected to be negative, or 1 when positive (SM Methods). Based on the signed graphs, we merged pairs of regime-shifts networks and created three response variables matrices that correspond to each of the cascading effects (Fig 1).

**Figure 1:**
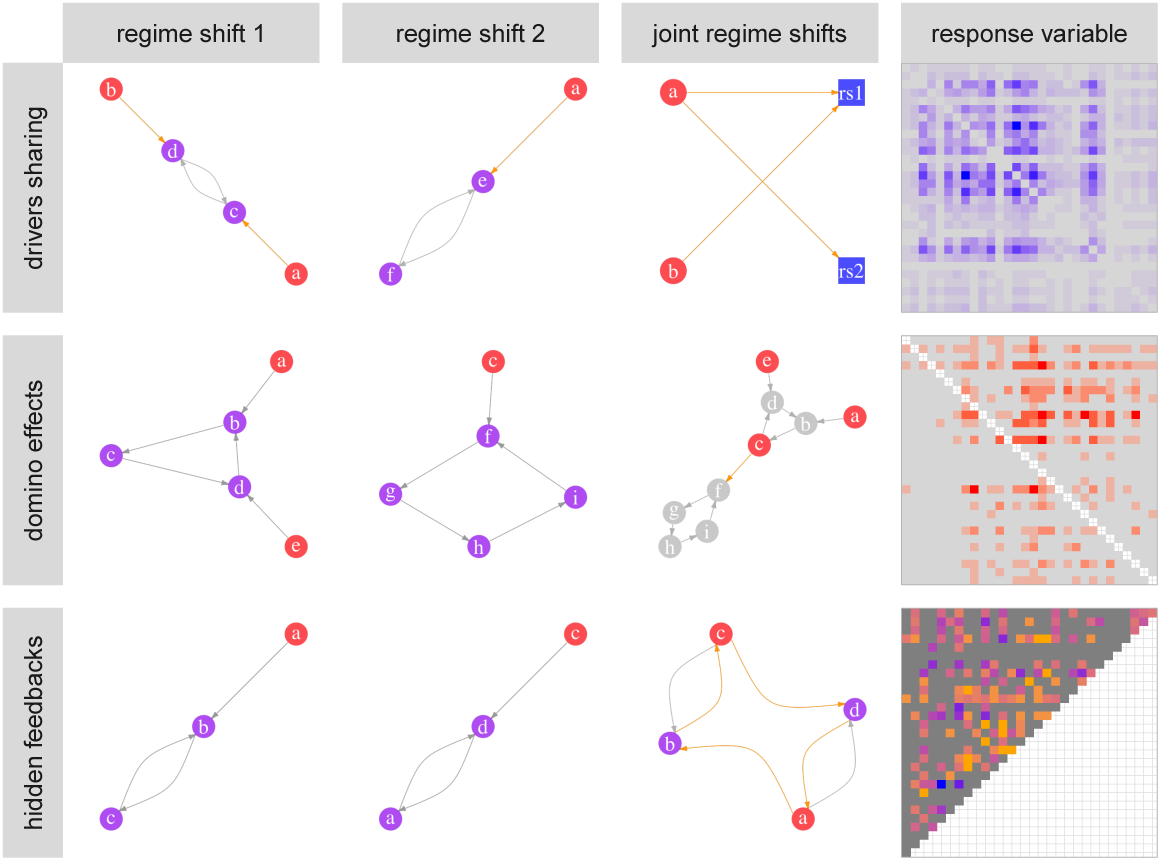
Minimal examples of the cascading effects. We merged pairs of regime shifts causal networks and created a response variable matrix that accounted for drivers shared, domino effects, or hidden feedbacks for all pair-wise combinations of regime shifts. For example, for drivers sharing two minimal regime shifts are depicted as causal diagrams, drivers are coloured red and variables inside feedbacks purple. The joint network is represented as a 2-mode network which allows us to study the co-occurrence of drivers across regime shifts. The response variable matrix counts the drivers shared by all pair-wise combinations of regime shifts. In the example of a domino effect two regime shifts are joined together where driver c in regime shift 2 is part of a feedback process in regime shift 1, creating a one-way dependency (orange colored link) between the two regime shifts. The response variable matrix counts all the one-way causal pathways between pair-wise combinations of regime shifts. In the example of a hidden feedback, two minimal regime shifts when joined together give rise to a new unidentified feedback of length 4 (orange circular pathway). The response variable matrix counts all hidden feedbacks that arise when merging all pair-wise combinations of regime shifts. Labelled matrices of the resulting response variables are available in the supplementary material.

We tested our three hypotheses using as response variable for *drivers sharing* the number of drivers shared, for *domino effects* the number of directed pathways that connect two regime shifts, and for *hidden feedbacks* it is the number of k-cycles (where *k* denotes cycle length) that emerge on the joined network that do not exist on the separate causal networks (Fig. 1, SM Methods). Each of these response variables are matrices that can be represented as weighted networks in our statistical framework, in which nodes are regime shifts and the link depends on the cascading effect described (Fig. 1). Thus, our research question can be rephrased as ‘What is the likelihood of a link between regime shifts in the response variable network, and what features change this likelihood?’ As explanatory variables we use the regime-shift database categorical variables (*N* = 75, Table S2, see SM methods for processing of variables). We focused on how similar two regime shifts are based on the categorical variables they share, and how the similarity increases or not the likelihood of having a link. We analyzed patterns of connections using exponential random graph models (SM Method). All these networks contain weighted links of count data; therefore, the specification for the models have a Poisson reference distribution (*23*).

### Driver sharing

Aquatic regime shifts tend to have and share more drivers, although the driver sharing is not exclusively with other aquatic regime shifts (Fig. 2a, Fig S3). The resulting matrix for *driver sharing* is a one-mode projection of the 2-mode network composed by 79 drivers and 30 regime shifts (Fig 1). The highest concentration of driver sharing is found between regime shifts in kelps, marine euthrophication, the collapse of fisheries and hypoxia. Terrestrial and polar regime shifts tend to have fewer and more idiosyncratic sets of drivers.

**Figure 2:**
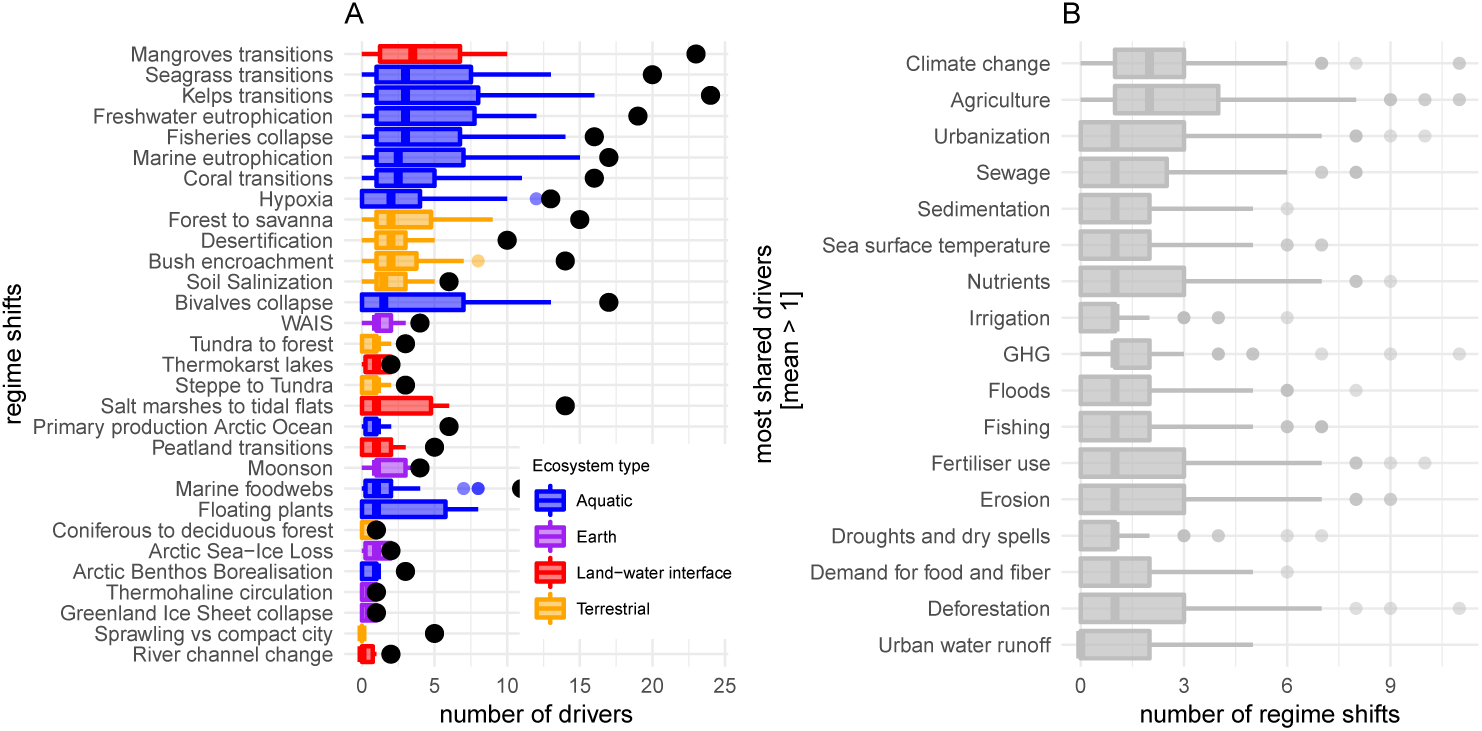
Driver sharing. The distribution of drivers shared per regime shift (a) with respect to the number of drivers each one has (black points) shows that regime shifts in aquatic environments tend to have and share more drivers. (b) shows the drivers most shared. WAIS abbreviates West Antarctica Ice Sheet collapse.

Large-scale regime shifts in polar and sub-continental areas (e.g. Monsoon weakening) have fewer drivers but are hotspots of sharing, typically including climate-related drivers. In support of previous results (*13*), we find that the most co-occurring drivers are related to food production, climate change and urbanization (Fig. 2b). Regime shifts are more likely to share drivers when they occur in similar ecosystem types and impact similar regulating services (p << 0.001, Fig 3, Table S1). The likelihood of sharing drivers is also affected by occurring on similar land use, impacting similar cultural services and aspects of human wellbeing, matching spatial scales (but not temporal ones), as well as having similar types of evidence and reversibility (p < 0.05, Fig 3, see Table S3 for comparison with null and alternative models fitted).

**Figure 3:**
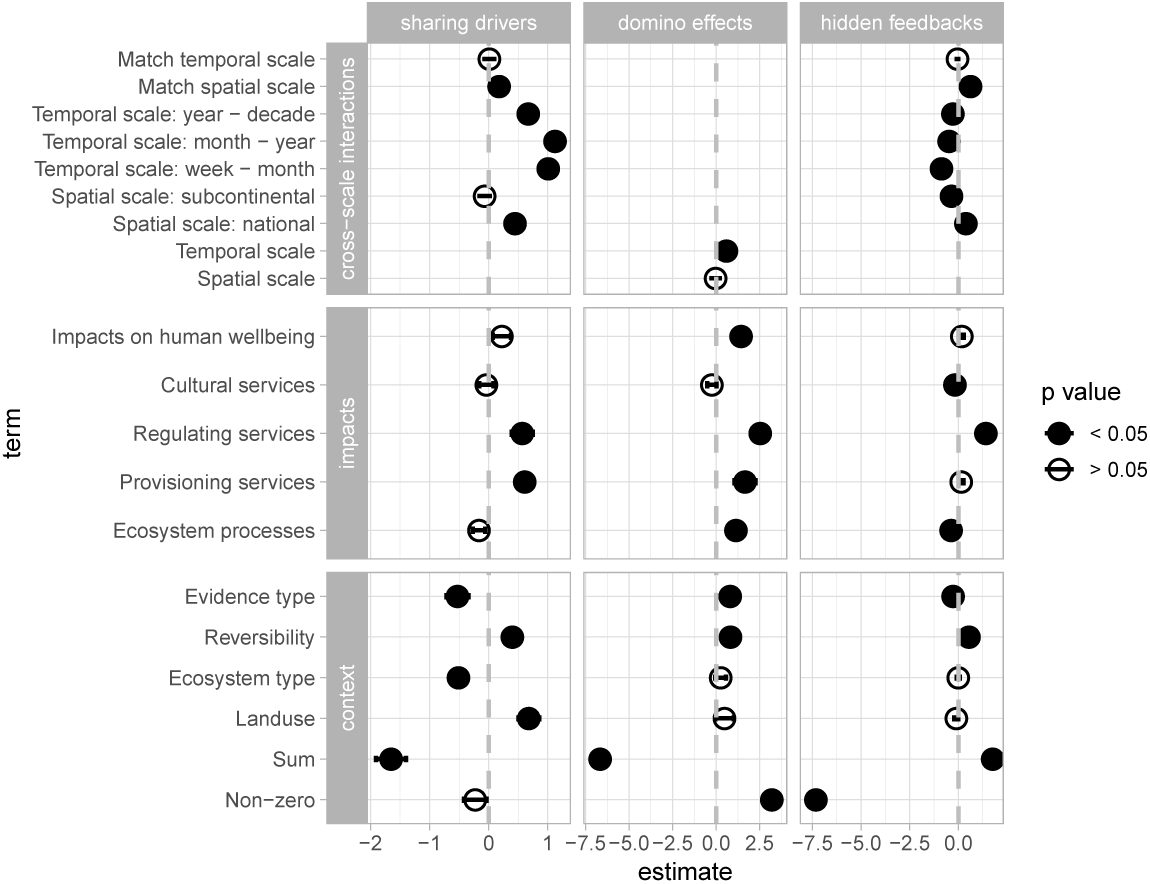
Statistical models summary. The response variables are the matrices of regime shifts interconnections for each cascading effect type (Fig 1). The terms fitted correspond to the categorical variables from the regime-shifts database (Table S2) that capture the similarity of regime shifts regarding their impacts on ecosystem services and human well being, the scales at which they occur, reversibility and evidence type (See SM Methods). Results from exponential random graph models are interpreted similarly to those of logistic regression where the non-zero and sum terms are the equivalent of an intercept. The non-zero term is the likelihood of the existence of a link in the response variable network, while the sum corresponds to its weight. Only best models are shown (lowest AIC or BIC), for complete results of null and alternative models fitted see Tables S3, S4 and S5.

Intuitively, the sharing of drivers is proposed as a potential mechanism that can correlate regime shifts in space and time, but not necessarily make them interdependent (*13*, *17*) unless they also share feedback mechanisms. Only 35% of all pair-wise combinations of regime shifts are coupled solely by driver sharing. Time correlation is debated given that spatial heterogeneity can break the synchrony induced by the sharing of drivers (*17*), meaning that contextual settings matter for such correlations to emerge. Spatial heterogeneity is also attributed as a mechanism that can smooth out critical transitions and soften their abruptness (*24*, *25*). Yet, with or without masking mechanisms, identifying common drivers is useful for designing management strategies that target bundles of drivers instead of well-studied variables independently, increasing the chances that managers will avoid several regime shifts under the influence of the same sets of drivers (*13*, *26*). For example, management options for drivers such as sedimentation, nutrient leakage, and fishing can reduce the likelihood of regime shifts such as eutrophication and hypoxia in coastal brackish lagoons, as well as coral transitions in adjacent coral reefs.

### Domino effects

Domino effects were investigated by searching variables that belong to feedback mechanisms in one regime shift and at the same time are drivers of another (Fig 1, see SM Methods). Despite having relatively fewer drivers, regime shifts that account for most domino effects include the Greenland ice sheet collapse, the Monsoon weakening, primary production in the Arctic Ocean, river-channel changes, soil salinization, weakening of the thermohaline circulation and the shift from tundra to forests (Fig 4). Conversely, the regime shifts that receive most influence through domino effects are mangrove transitions, kelp transitions and transitions from salt marshes to tidal flats. The maximum number of pathways found was 4, and the variables that produce most domino effects relate to climate, nutrients and water transport (Fig 4). In line with our expectations, regime shifts that contain variables that will in turn be drivers of other regime shifts typically have large spatial scales and slow temporal scales: thermohaline circulation collapse, river channel change, monsoon weakening, and Greenland ice sheet collapse. On the other hand, regime shifts that receive the influence are often marine and their time and space dynamics contained more locally. The statistical models support this observation (Fig 3): regime shifts whose time scales are on the range of weeks to months are more likely to receive influence from regime shifts whose dynamics occur on the scale of years to decades (p < 0.1), but we did not find evidence for spatial scales (Table S4). Both, having a link and having a high number of pathways are significant (p < 0.01). The odds of having higher numbers of domino effects are increased when regime shifts impact similar regulating services and similar aspects of human well-being (p < 0.001).

**Figure 4:**
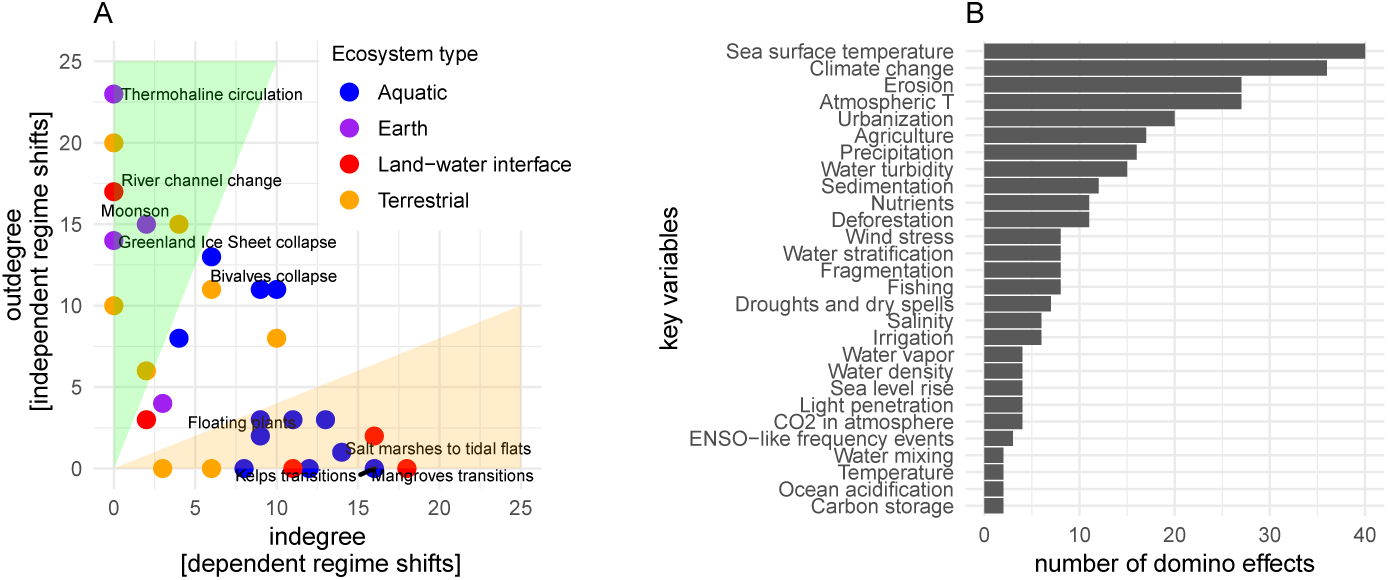
Domino effects. Regime shifts that produce most domino effects (high outdegree, shaded green area) are Earth system tipping points, while the regime shifts that receive the most (high indegree, orange shaded area) occur in aquatic and land-water interface (a); labels are plotted only for regime shifts where the maximum number of domino effects is found (4). Most variables associated with domino effects are related to climate and transport mechanisms (b). These variables are part of a feedback mechanism in one regime shift that are in turn drivers in another regime shift.

### Hidden feedbacks

Contrary to the results found for driver sharing, most hidden feedbacks occur in terrestrial and earth systems (Fig 5) and typically have higher feedback length. The regime shifts with higher numbers of connections (16 to 18 out of 30 possible) are forest to savanna, monsoon weakening, thermohaline circulation, desertification, primary productivity of the Arctic Ocean and the Greenland ice sheet collapse. Key variables that belong to many of these hidden feedbacks are related to climate, fires, erosion, the functional role of herbivores, agriculture and urbanisation (Fig 5). The statistical analysis (Fig 3) shows that there is fewer hidden feedbacks that one would expect by random, but when they do occur, the odds of having multiple feedbacks coupling two regime shifts is 7.33 times higher. The odds of two regime shifts been connected through hidden feedbacks is not affected by occurring on similar land uses or ecosystem types. Yet, the likelihood of hidden feedbacks increases if the pair of regime shifts impact similar ecosystem processes, impact similar regulating and cultural services, and especially if they match spatial scales but not necessarily temporal ones (Fig 3, p < 0.001, see table S5 for null and alternative models fitted), suggesting cross-scale interactions in time.

**Figure 5:**
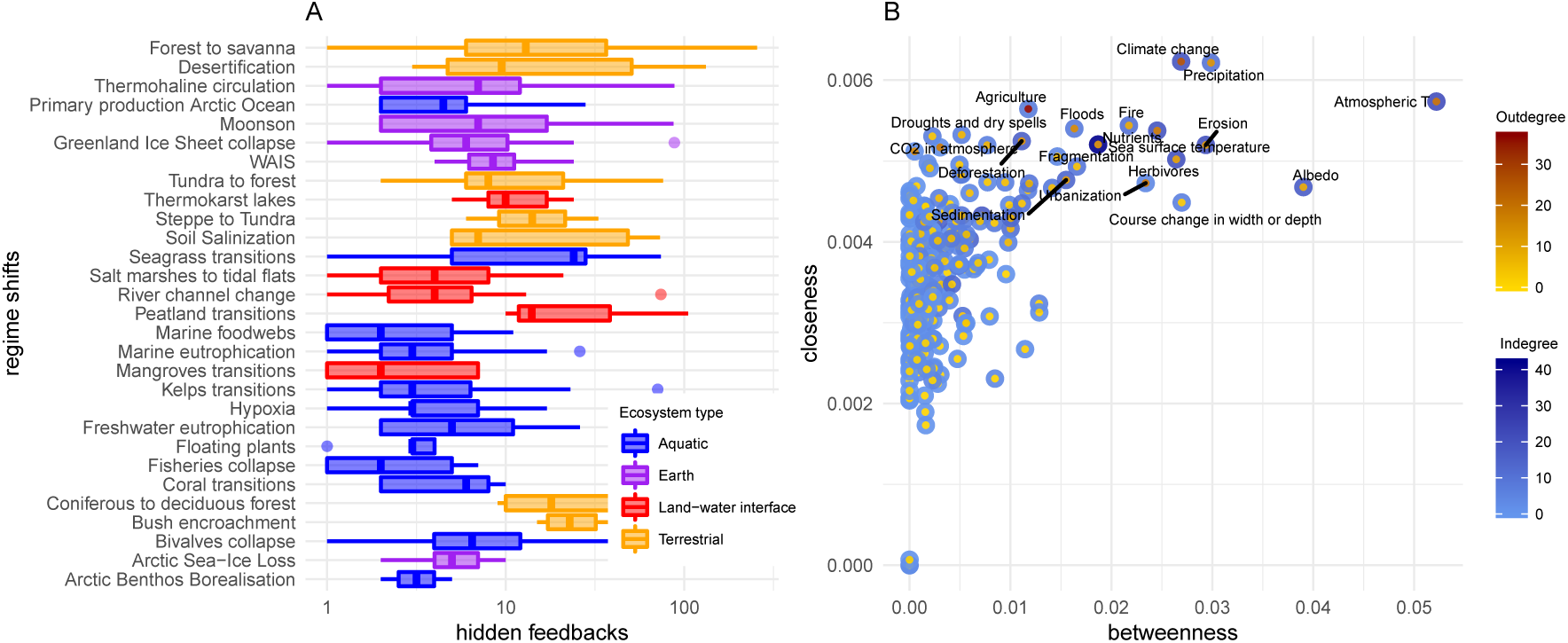
Hidden feedbacks. Hidden feedbacks occur typically in terrestrial and Earth system regime shifts, (a) shows the distributions of hidden feedbacks and has been organised by higher to lower mean in the number of feedbacks. Boxplots are shown in log-scale after zero values have been removed. The variables most often involved in hidden feedbacks have high betweenness and closeness centralities (b) calculated on the network of all regime shifts in our sample (N = 30). These measures reveal the variables (labelled) that lie on most shorter pathways from all other variables in the network

Out of the 870 pair-wise combinations of regime shifts analysed (when taking into account directionality), ~35% are solely coupled through sharing drivers, ~4% through domino effects, and ~2% through hidden feedbacks (Fig 6). Furthermore, 28% of these pair-wise combinations are coupled through two different types of connections, and 10% by all three of them. Only for 167 (19%) pair-wise combinations we can be certain with the current dataset that there is no cascading effects. However, the discovery of new drivers or feedback mechanisms underlying these dynamics can reduce this estimate.

**Figure 6:**
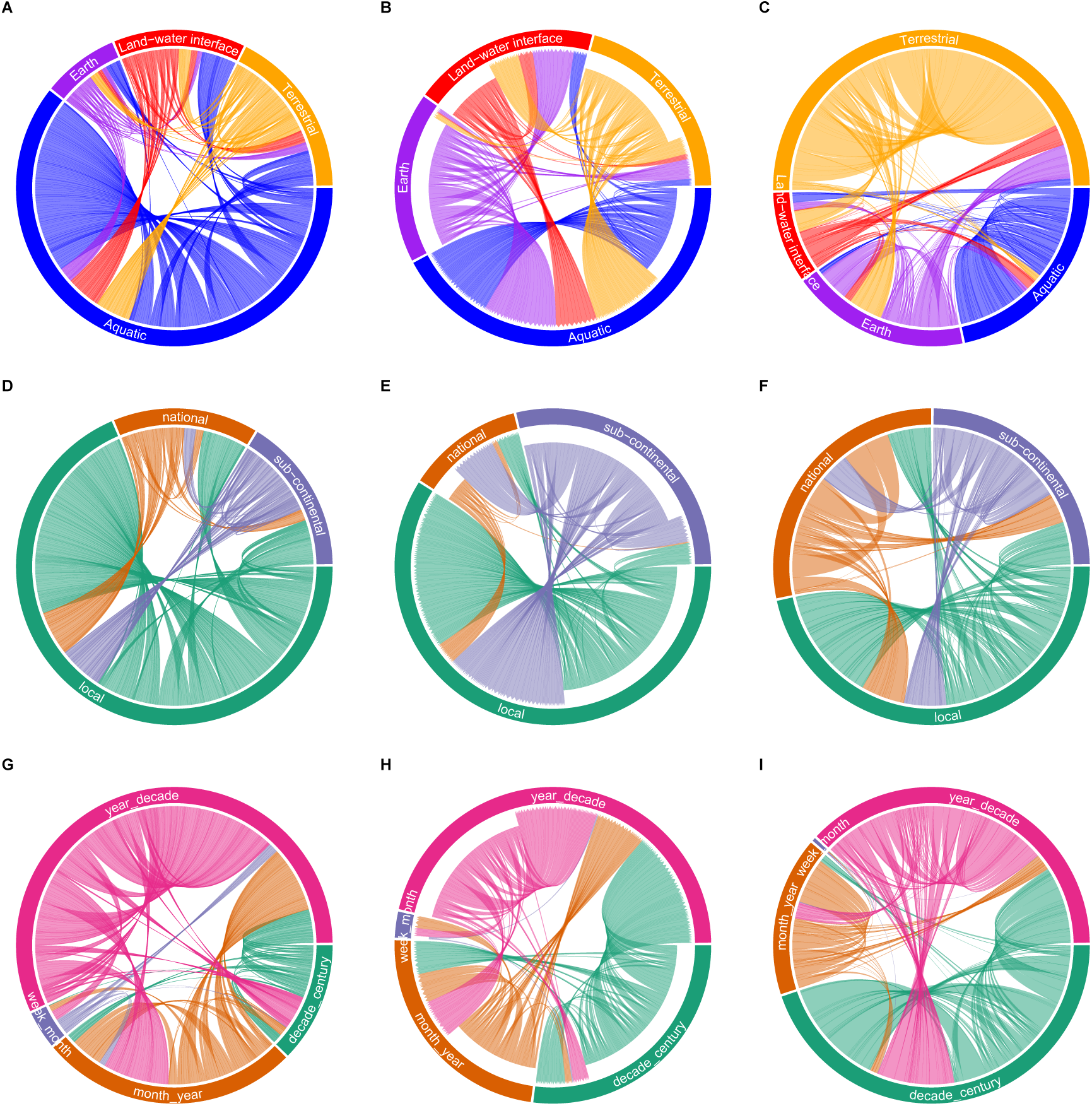
Summary of cascading effects and cross-scale interactions. Networks of regime shifts interactions are shown by cascading effect type: sharing drivers (a,d,g), domino effects (b,e,h), and hidden feedbacks (c,f,i). These circular maps show the links between regime shifts found classified by regime shift type (a,b,c), spatial scales (d,e,f), and temporal scales (g,h,i). Link width is scaled according to the number of shared drivers, domino effects or hidden feedbacks found. Figure S3 complements this figure by showing the matrix where multiple connections are expected as well as the pair-wise combinations where no connections were found.

**Figure 7:**
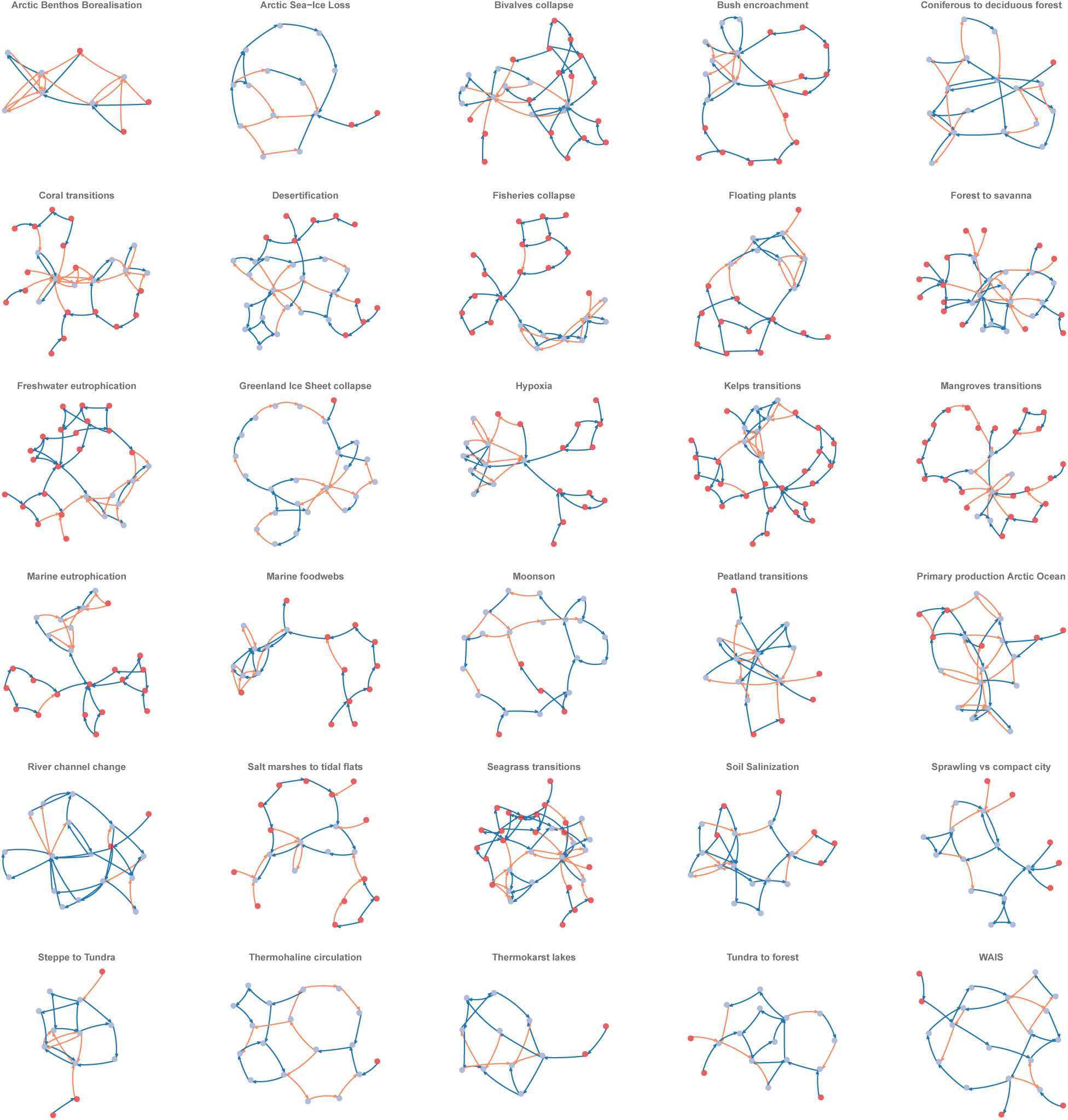
Fig S1. Causal networks for all regime shifts analysed

**Figure 8:**
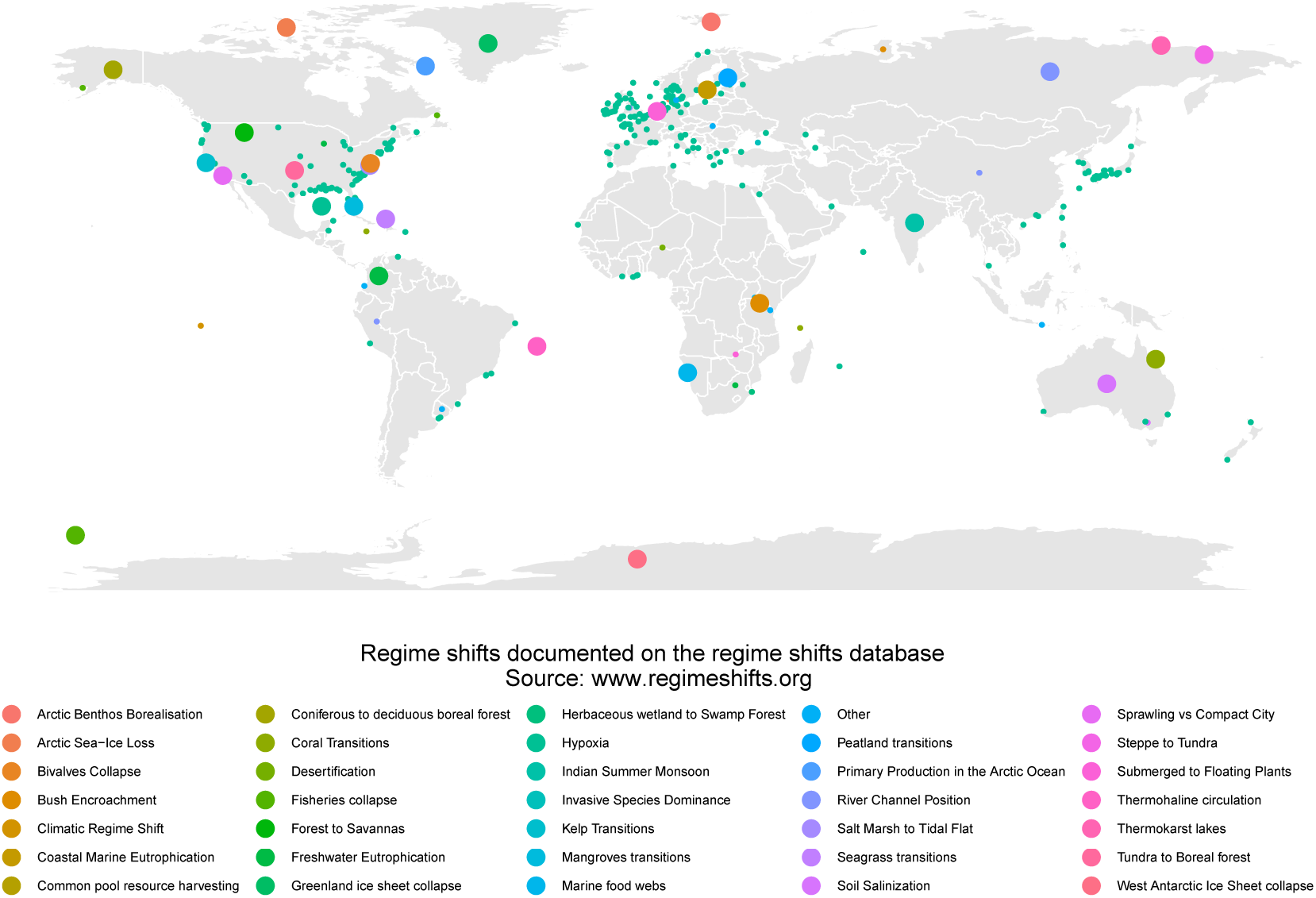
Fig S2. Regime shifts around the world. Large points show generic types of regime shifts (n = 35) while small points are case studies (n=324).

**Figure 9:**
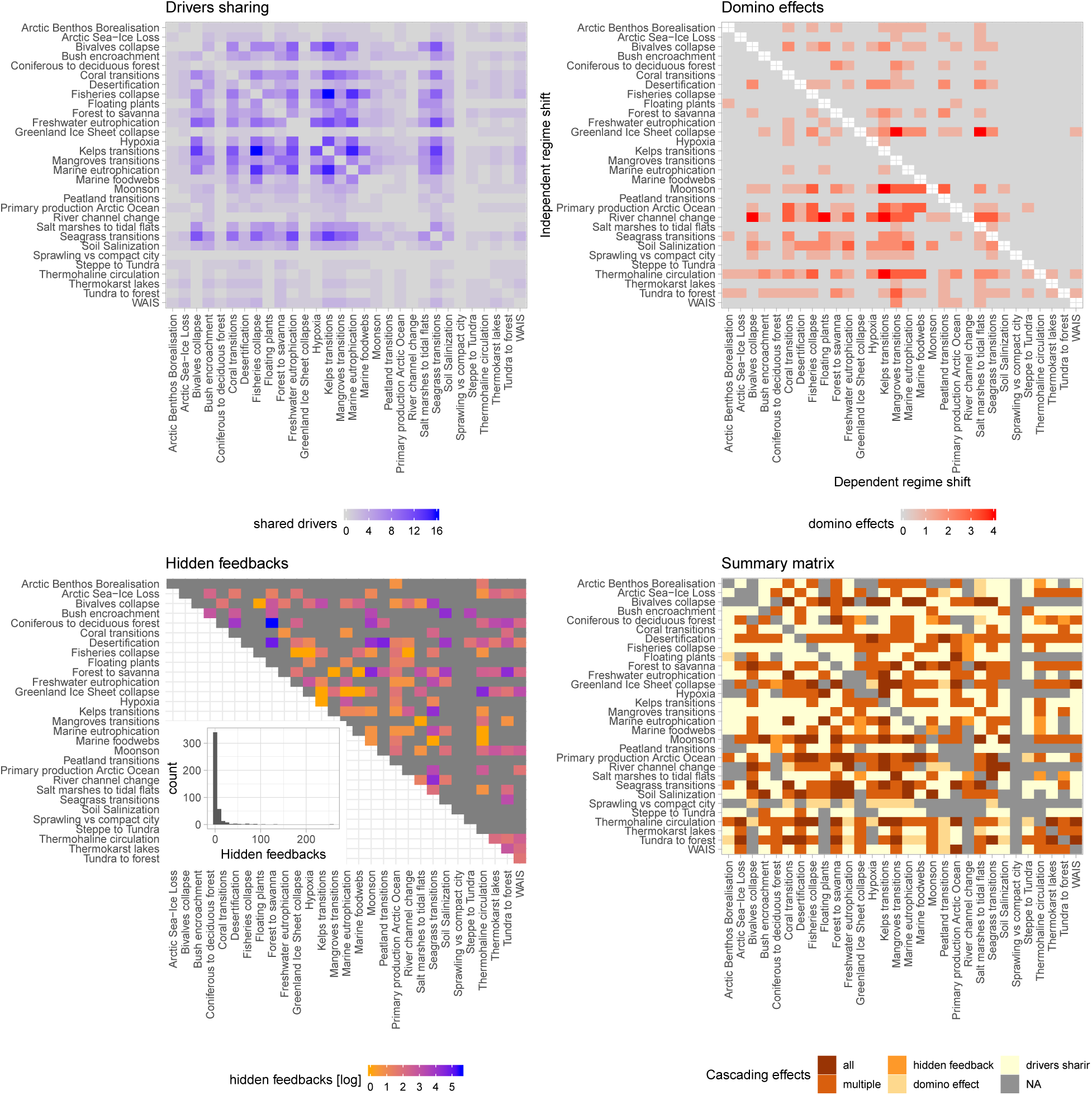
Response variable matrices and summary of all cascading effects in matrix form.

## Discussion

Our results show we can expect ecosystems to experience cascading regime shifts. Regime shifts that occur in similar ecosystem types and land uses are more likely to share drivers, confirming our expectation. Our analysis also suggests that sharing drivers increase when regime shifts operate at similar spatial scales, or when they impact similar provisioning or regulating services. We expected that domino effects will be more common between regime shifts whose dynamics occur at larger spatial scales and slower dynamics in time towards regime shifts more localised in space and faster in time. However, we only found support for the cross-scale dynamic in time but not in space. Additionally, aquatic regime shifts often received the domino effect supporting the idea that water acts as a potential transport mechanism linking regime shifts in one-way interactions (*27*). Last, we expected that hidden feedbacks would emerge when scales match both in space and time, as well as under similar ecosystem types. Our results only confirm matching in spatial scales and also suggest that hidden feedbacks are common when regime shifts impact similar ecosystem processes, regulating and cultural ecosystem services.

We found 45% of all pair-wise combinations of regime shifts are connected by a plausible domino effect or hidden feedbacks (Fig 6, Fig S3). Domino effects and hidden feedbacks are often disregarded because research on regime shifts is divided by disciplines and tipycally focus on one system at the time. Consequently, data collection and hypothesis testing for coupled systems has largely remained unexplored (*16*, *17*). Our findings align with previous results on the type of variables and processes that might couple distant regime shifts (See Table S1), highlighting the role of climate and transport mechanisms for nutrients and water (Figs 2, 4, 5).

Recent literature (refs. in Table S1) reports potential linkages between euthrophication and hypoxia, hypoxia and coral transitions, shifts in coral reefs and mangroves transitions, or climate interactions. Other examples in the terrestrial realm report potential increase in Arctic warming from higher fire frequency in boreal forest or permafrost thawing. Regime shifts in the Arctic can impact any temperature driven regime shift in and outside the Arctic (*28*), including the weakening of the thermohaline circulation. Moisture recycling is a key underlying feedback on the shift from forest to savanna or the Indian monsoon; but also has the potential to couple ecosystems beyond the forest that depend on moisture recycling as an important water source. Changes in moisture recycling can affect mountain forest in the Andes, nutrient cycling in the ocean by affecting sea surface temperature and therefore regime shifts in marine food webs, or exacerbation of dry land related regime shifts. These connections between regime shifts represent an emergent literature on regime shifts interactions (Table S1 and references therein). We contribute to this endeavour with a network-based method that allow us to explore plausible cascading effects and distinguish potential correlations from true interdependencies. Distinguishing whether the coupling is expected to be correlational because of driver sharing, a one-way causation (the *domino effect*), or a two-way interaction (the *hidden feedbacks*), provides a useful set of hypotheses for future research.

While our method identifies plausible connections between regime shifts, identifying under what conditions plausible becomes probable requires more detailed understanding on regime shifts mechanisms. Empirical studies and modelling syntheses are required to translate our identification of possible mechanisms into context sensitive forecasts. Dynamic models of this type of dynamics require careful assumptions about parameter values as well as functional form of the system’s equations. Generalised modelling is a promising technique that does not require particular assumptions allowing the researcher to reach more general conclusions based on stability properties of the system (*29*, *30*). Another potential avenue for future research is looking at how transport mechanisms couple far-apart ecosystems. One example already mentioned is the moisture recycling feedback (*31*). Another important social teleconnection (sensu (*16*)) could be with allocation of resources through international trade, investigating how demand of resources in certain countries can shape the state space of ecosystems from the providing countries. An important lesson from our study is that regime shifts can be interconnected, they should not be study in isolation assuming they are independent systems. On the contrary, methods and data collection that takes into account the possibility of cascading effects needs to be further developed.

The frequency and diversity of regime shifts interconnections suggests that current approaches to environmental management and governance are substantially underestimating the likelihood of cascading effects. More attention should be paid to how the Earth is social-ecologically connected (*16*), how those connections should be managed, and how to best prepare for regime shifts. Our research suggests that regional ecosystems can be transformed by ecosystem management far away, and conversely, can themselves drive the transformations of other distant ecosystems. Decisions made in one place can undermine the achievement of sustainable development goals in other places. For example, it has been shown that what happens in the Arctic does not stay in the Arctic (*28*, *32*), many Arctic regime shifts have the potential to impact non-Arctic ecosystems far away and the provision of their ecosystem services. It implies that whoever does make decisions on management is not necessarily the one that has to deal with the impacts. This issue is evident in governance of water transport systems, whether run-off or atmospheric transport, but it is applicable to other dynamics connecting far-away ecosystems through other mechanisms such as climate change, fire, nutrient inputs or trade. Our results highlight variables that are key for domino effects and hidden feedbacks. They are also good observables for monitoring early warning indicators of the strengthening of regime-shifts coupling.

## Conclusions

Regime shifts occur across a wide range of ecosystems. However, how a regime shift somewhere in the world could affect the occurrence of another regime shift remains an open question and a key frontier of research. We propose two types of cascading effects that can connect different regime shifts: domino effects, and hidden feedbacks; and compare them with the case of driver sharing. To assess these cascading effects among regime shifts we developed a network-based method that identified plausible cascading effects. Regime shifts that occur in similar ecosystems and land uses are more likely to share drivers, and consequently respond similarly to environmental change. One-way directional interactions or *domino effects* are common but not exclusive to aquatic regime shifts, it highlights the role of water transport and tend to emerge between regime shifts whose dynamics occur at similar temporal scales. Two-way interconnections or *hidden feedbacks* are more common in terrestrial systems, they occur often between regime shifts that match spatial scales and impact similar ecosystem processes. These results suggest that the Anthropocene can be expected to increase ecological surprise. How and when nonlinear change can be transmitted across space and time in the earth system should be considered in assessments of future environmental change and planning.

## Acknowledgements

We are grateful to the contributors, reviewers and developers of the regime shifts database. This work was supported by FORMAS grant 942-2015-731 to JR. JR designed the research, JR and GP curated the data, JR wrote the code and run the analysis with guidance from GP, ÖB and SL; JR, GP, ÖB and SL wrote the paper. Authors declare no competing interests. Data from the regime shifts database is publicly available at www.regimeshifts.org. Upon publication, the version of the database used, curated causal networks, and code will be made available in both the regime shifts database and an online repository (e.g. figshare or dryad). The development version of the code is available at: https://github.com/juanrocha/Domino

## Supplementary material

**Table 1:** Summary of examples from the literature

| Regime shift 1 | Regime Shift 2 | Type | Reference |
| --- | --- | --- | --- |
| Marine eutrophication | Hypoxia | Sharing drivers,<br>domino effect | (14) |
| Hypoxia | Coral transitions | Domino effect | (15) |
| Forest to savanna | Mountain forest<br>transitions | Domino effect | (33, 34) |
| Forest to savanna | Marine food webs | Domino effect | (35) |
| Forest to savanna | Dryland degradation | Domino effect | (36) |
| Coniferous to<br>deciduous boreal forest | Arctic sea ice loss | Hidden feedback | (37, 38) |
| Peatland transitions | Arctic sea ice loss | Hidden feedback | (39) |
| Arctic sea ice loss | Thermohaline<br>circulation weakening | Hidden feedback | (40) |
| Antarctic sea ice loss | Marine food webs | Hidden feedback | (41) |
| Thermohaline<br>circulation weakening | ENSO | Domino effect | (22) |

### Supplementary methods

This section complements the methods and results from the exponential random graph models. The modeling technique used the number of drivers shared, the number of domino effects and the number of hidden feedback as the three key response variables (Fig 1). The use of exponential random graph models is motivated because it take into account network structure and allow us to control for sample bias. It investigates the odds of the existence and weight of a link and its results are interpreted similarly to a logistic regression.

### Data

The regime shifts database synthesizes scientific literature on our current knowledge on regime shifts. It is an open online repository of regime shift syntheses and case studies. When possible, entries to the regime shifts database have been peer-reviewed by an expert on the topic to ensure quality and accuracy of its contents (*10*). The main data used in this study are the causal loop diagrams described in the database (Fig S1), and a set of categorical variables about regime shifts attributes that were used to fit statistical models to explore the role of cross scale interactions (described below). Out of the 35 regime shifts documented in the database, here we use 30 where complete synthesis exist and causal loop diagrams have been curated.

*Causal loop diagrams* (CLDs) represent a collection of causal mechanisms that scientist have reported in their narratives: both empirical (what they choose to sample) or theoretical (what they choose to model). CLDs consist of variables connected by arrows denoting causal influence (*42*), also known as signed directed graphs. Each relationship must have a positive (+) or negative (–) sign that represents the effect of the dependent variable given change on the independent variable (*42*, *43*). Although the functional form that underlies each relationship is not necessarily known, positive relationships are proportional while negative ones are inversely proportional *ceteris paribus.* CLDs assume that the causal relationships captured by links are monotonic, while non-linearities are captured by links sets: thus they do not have self-loops. Feedback loops are the basic structural units of the diagram and emerge by connecting variables in closed directed paths (cycles). Feedback means that once a signal enters the loop, some part of the output is fed back to the input, resulting in amplification or dampening of its own signal. Feedbacks can be reinforcing if the overall polarity of its links is positive, or balancing if negative. Reinforcing feedbacks are usually responsible for behaviors that drive the system out of equilibrium, while balancing feedbacks are responsible for near equilibrium dynamics such as oscillations and delays (*42*). Note that causal links do not describe the behavior of variables, only the structure of the system: they describe what would happen if there were changes (*42*). CLDs were curated in the regime shifts database in a way that variables names are consistent (e.g. agriculture and cropping is kept as ‘agriculture’), and feedback loops comparable (e.g. albedo in the rainforest, Arctic or Antarctic regime shifts is the same feedback). Fig S1 shows all CLD’s used in the analysis.

### Categorical variables

We calculated the similarity of each pair-wise combination of regime shifts in the database regarding categorical attributes (Table S2) such as *(i)* land use under which the regime shift occur, *(ii)* ecosystem type, impacts on *(iii)* ecosystem processes, *(iv)* provisioning services, *(v)* regulating services, *(vi)* cultural services, *(vii)* and human wellbeing; as well as *(viii)* the spatial scale at which the regime shift occur, *(ix)* the temporal scales, *(x)* reversibility, and *(xi)* evidence type (*10*, *13*). For all 75 categorical variables encoded, the database reports presence or absence (0, 1) allowing us to calculate the Jaccard index and use it as a proxy of how similar two regime shifts are. To facilitate the interpretation of statistical models, the Jaccard distance was rescaled (*x* = 1 – *J_d_*) so equivalent regime shifts score 1 and complete dissimilar zero. Given that most of our hypotheses relate to the scale at which regime shifts occur, we also modelled scale as a categorical variable to account for matches in the network, not only similarity. Figure S2 shows a map with all generic types of regime shifts and case studies reported in the regime shifts database.

**Table 2:**
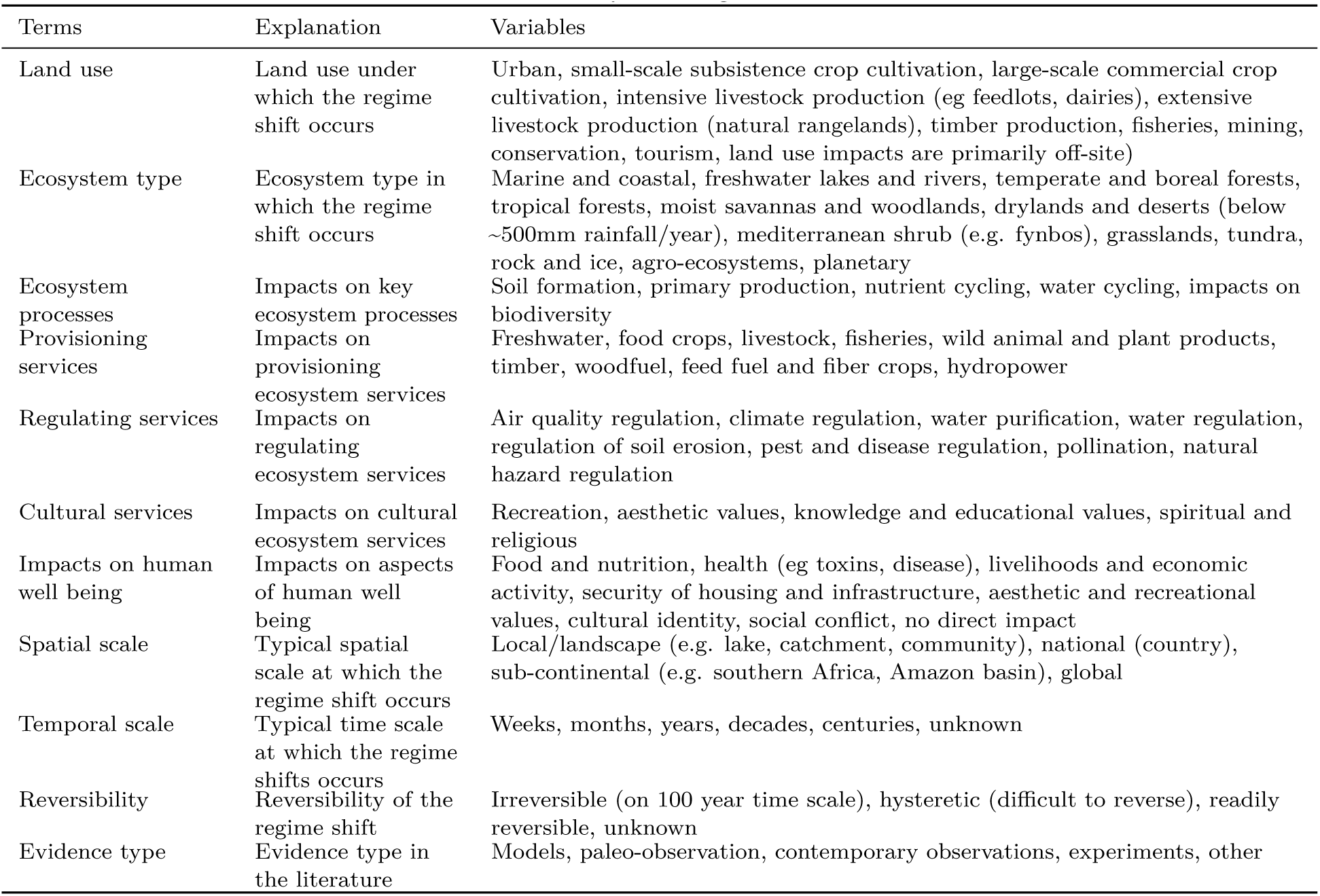
Summary of categorical variables

| Terms | Explanation | Variables |
| --- | --- | --- |
| Land use | Land use under which the regime shift occurs | Urban, small-scale subsistence crop cultivation, large-scale commercial crop cultivation, intensive livestock production (eg feedlots, dairies), extensive livestock production (natural rangelands), timber production, fisheries, mining, conservation, tourism, land use impacts are primarily off-site) |
| Ecosystem type | Ecosystem type in which the regime shift occurs | Marine and coastal, freshwater lakes and rivers, temperate and boreal forests, tropical forests, moist savannas and woodlands, drylands and deserts (below ~500mm rainfall/year), mediterranean shrub (e.g. fynbos), grasslands, tundra, rock and ice, agro-ecosystems, planetary |
| Ecosystem processes | Impacts on key ecosystem processes | Soil formation, primary production, nutrient cycling, water cycling, impacts on biodiversity |
| Provisioning services | Impacts on provisioning ecosystem services | Freshwater, food crops, livestock, fisheries, wild animal and plant products, timber, woodfuel, feed fuel and fiber crops, hydropower |
| Regulating services | Impacts on regulating ecosystem services | Air quality regulation, climate regulation, water purification, water regulation, regulation of soil erosion, pest and disease regulation, pollination, natural hazard regulation |
| Cultural services | Impacts on cultural ecosystem services | Recreation, aesthetic values, knowledge and educational values, spiritual and religious |
| Impacts on human well being | Impacts on aspects of human well being | Food and nutrition, health (eg toxins, disease), livelihoods and economic activity, security of housing and infrastructure, aesthetic and recreational values, cultural identity, social conflict, no direct impact |
| Spatial scale | Typical spatial scale at which the regime shift occurs | Local/landscape (e.g. lake, catchment, community), national (country), sub-continental (e.g. southern Africa, Amazon basin), global |
| Temporal scale | Typical time scale at which the regime shifts occurs | Weeks, months, years, decades, centuries, unknown |
| Reversibility | Reversibility of the regime shift | Irreversible (on 100 year time scale), hysteretic (difficult to reverse), readily reversible, unknown |
| Evidence type | Evidence type in the literature | Models, paleo-observation, contemporary observations, experiments, other |

### Networks

Causal loop diagrams were reinterpreted as networks where node attributes were coded if a node belonged to a feedback loop -a k-cycle in the network- or not. The later is then by definition a driver, an independent variable whose dynamics are not affected by the dynamics of the state variables of the system at hand. Note that networks in ecology usually describe inter-species interactions such as predation or mutualism. Our approach is different from what has been done in ecology. Here a network describes a set of processes, both biotic and abiotic, that can govern ecosystem regime-shift dynamics. Our approach is inspired by other network applications to processes such as cell metabolic networks or the network of human diseases, where a process is not captured by a link type (e.g. predation) but by a collection of link interactions (e.g. the Krebs cycle). Thus, individual species are not taken into consideration, but rather their functional roles at the aggregated ecosystem scale (e.g. herbivory).

### Shared drivers

To study shared drivers we created a bipartite network of drivers and regime shifts following the method outlined by Rocha et al (*13*). The statistical models were performed for the case of driver sharing on the one-mode network projection of regime shifts sharing drivers (this is the matrix *A^T^A*), and categorical variables from the regime-shifts database were used as node attributes or node covariates to test our hypothesis.

### Domino effects

occur when the occurrence of a regime shift can increase or decrease the likelihood of other regime shifts occurring, creating a one-way dependence. Different from the driver-sharing network, here we explore potential domino effects by using the full causal loop diagrams as networks. The algorithm for identifying domino effects takes the adjacency matrix of two given regime shifts *A*_1_ and *A*_2_ and identifies all nodes *n* ∈ *A*_1_ ∩ *A*_2_ such that *n* belong to a feedback in *A*_1_ but is a driver in *A*_2_. Thus, set differences between causal pathways suggest missing drivers, and set intersection between causal pathways and feedback loop nodes indicate potential domino effects (Fig. 2b). By iterating this simple algorithm we derived how many different pathways exist between every pair-wise combination of regime shifts (*N* = 870). The resulting non-symmetrical matrix represents a directed network with regime shifts as nodes and link weights as the number of pathways used for statistical analysis.

### Hidden feedbacks

We explore hidden feedbacks by pair-wise comparison of causal networks. First, the feedback loops (or k-cycles) are counted by feedback length *k* for each regime shift matrix *A*_1_ and *A*_2_ separately. Then the cycle count is applied to the composite network *A*_1,2_ of the two regime shifts. The difference between the k-cycles in the composite network *A*_1,2_ and the k-cycles in the individual networks *A*_1_ and *A*_2_ are the hidden feedbacks that emerge when the two causal networks are joined. By iterating the same procedure to all pair-wise combinations of regime shifts (*N* = 435) we obtain a symmetric directed matrix that is then used for the statistical analysis. Although computationally intensive, the search for cycles is feasible in our networks given the small size and relative sparse structures. In fact, the maximum cycle length *k* is bounded by the size of the network. Out of the 435 coupled networks analysed, the maximum feedback length was 56.

### Exponential random graph models

The coefficients in exponential random graph models present the log-odds of the existence of a link given the similarites (Jaccard distances) or matching of node attributes (e.g. scale). Thus, their results are read similarly to those of a logistic regression. Instead of the intercept here we have the `nonzero` term which indicates what the odds are of the existence of a link. Since the networks are weighted by the number of shared drivers, domino effects or hidden feedbacks, the term `sum` accounts for the weight of the link when it exists. We used Jaccard similarity on the attributes coded in the regime shifts database to quantify how similar two regime shifts are and how the similarity affects the odds of a link. Since our hypotheses were related to the temporal and spatial scales at which the regime shifts occurred, we also modeled such terms not only as similarity (a score between 0 and 1), but also as categorical mutually exclusive variables. The original data from the regime shifts database coded for categorical non-exclusive variables. For spatial scale, when two different categories where present (e.g. local, national), we choose the larger category as the unique factor. For temporal scales, regime shifts were often coded as belonging to two categories (e.g. decades and centuries), thus we create a variable `time-range` that bundles both minimum and maximum reported. Spatial and temporal scales are then also modelled as factors too, the term `nodefactor` in each model accounts for how more likely is to have a link when the factor is present, while the `nodematch` term accounts for the homogeneity effect of two nodes sharing the same node attribute (same scale). The analysis was performed in R statistical language (*44*) using packages for modeling networks with exponential random graph models (*45*–*48*).

**Table S3.**
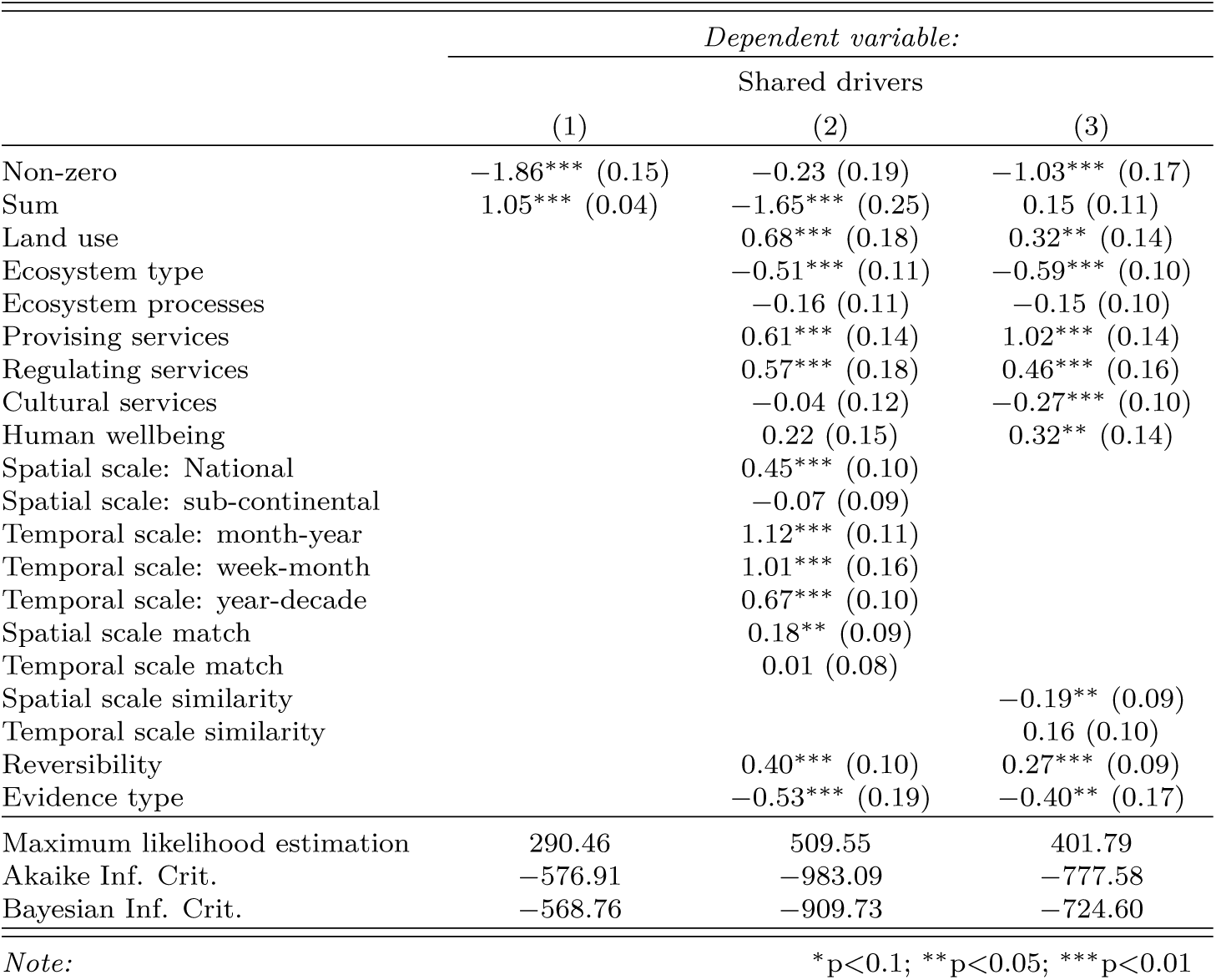
Exponential random graph models for *shared drivers*

**Table S4.** Exponential random graph models for *domino effects*

|  | *Dependent variable:* |  |  |  |
| --- | --- | --- | --- | --- |
|  | Domino effects |  |  |  |
|  | (1) | (2) | (3) | (4) |
| Non-zero | −1.63*** (0.16) | 3.20*** (0.36) | 3.33*** (0.38) | 3.44*** (0.39) |
| Sum | −0.06 (0.09) | −6.67*** (0.43) | −7.24*** (0.66) | −7.31*** (0.71) |
| Land use |  | 0.48 (0.53) | 0.70 (0.56) | 0.77 (0.58) |
| Ecosystem type |  | 0.26 (0.31) | 0.18 (0.32) | 0.31 (0.32) |
| Ecosystem processes |  | 1.14*** (0.35) | 1.03*** (0.38) | 0.82** (0.38) |
| Provising services |  | 1.66*** (0.60) | 1.92*** (0.64) | 1.97*** (0.65) |
| Regulating services |  | 2.53*** (0.42) | 2.69*** (0.44) | 2.87*** (0.45) |
| Cultural services |  | −0.25 (0.26) | −0.50* (0.27) | −0.64** (0.29) |
| Human wellbeing |  | 1.43*** (0.42) | 1.45*** (0.45) | 1.39*** (0.45) |
| Spatial scale similarity |  | −0.03 (0.23) |  |  |
| Temporal scale similarity |  | 0.59** (0.29) |  |  |
| Spatial scale: national |  |  | 0.04 (0.29) |  |
| Spatial scale: sub-continental |  |  | 0.27 (0.37) |  |
| Temporal scale: year |  |  | 0.11 (0.23) |  |
| Temporal scale: month |  |  | 0.22 (0.43) |  |
| Temporal scale: decade |  |  | −0.20 (0.19) |  |
| Indegree when spatial scale national |  |  |  | −0.16 (0.31) |
| Indegree when spatial scale subcontinental |  |  |  | −0.28 (0.56) |
| Indegree when temporal scale year |  |  |  | 0.08 (0.36) |
| Indegree when temporal scale month |  |  |  | 0.94* (0.52) |
| Indegree when temporal scale decade |  |  |  | 0.16 (0.30) |
| Outdegree when spatial scale national |  |  |  | −0.09 (0.60) |
| Outdegree when spatial scale subcontinental |  |  |  | 0.29 (0.48) |
| Outdegree when temporal scale year |  |  |  | 0.43 (0.38) |
| Outdegree when temporal scale month |  |  |  | −1.76 (1.37) |
| Outdegree when temporal scale decade |  |  |  | −0.48* (0.29) |
| Spatial scale match: local |  |  | 0.85** (0.41) | 0.60 (0.46) |
| Spatial scale match: national |  |  | 0.42 (0.89) | 0.36 (0.88) |
| Spatial scale match: sub-continental |  |  | 0.15 (0.48) | 0.48 (0.61) |
| Temporal scale match |  |  | 0.07 (0.19) | 0.20 (0.21) |
| Reversibility |  | 0.82*** (0.28) | 0.99*** (0.31) | 0.94*** (0.32) |
| Evidence type |  | 0.81** (0.41) | 1.38*** (0.48) | 1.28*** (0.48) |
| Maximum likelihood estimation | 279.89 | 676.49 | 683.9 | 691.54 |
| Akaike Inf. Crit. | −555.77 | −1,326.98 | −1,327.80 | −1,333.09 |
| Bayesian Inf. Crit. | −546.23 | −1,264.99 | −1,232.43 | −1,213.88 |

*Note:*
\*p<0.1; \*\*p<0.05; \*\*\*p<0.01

**Table S5.**
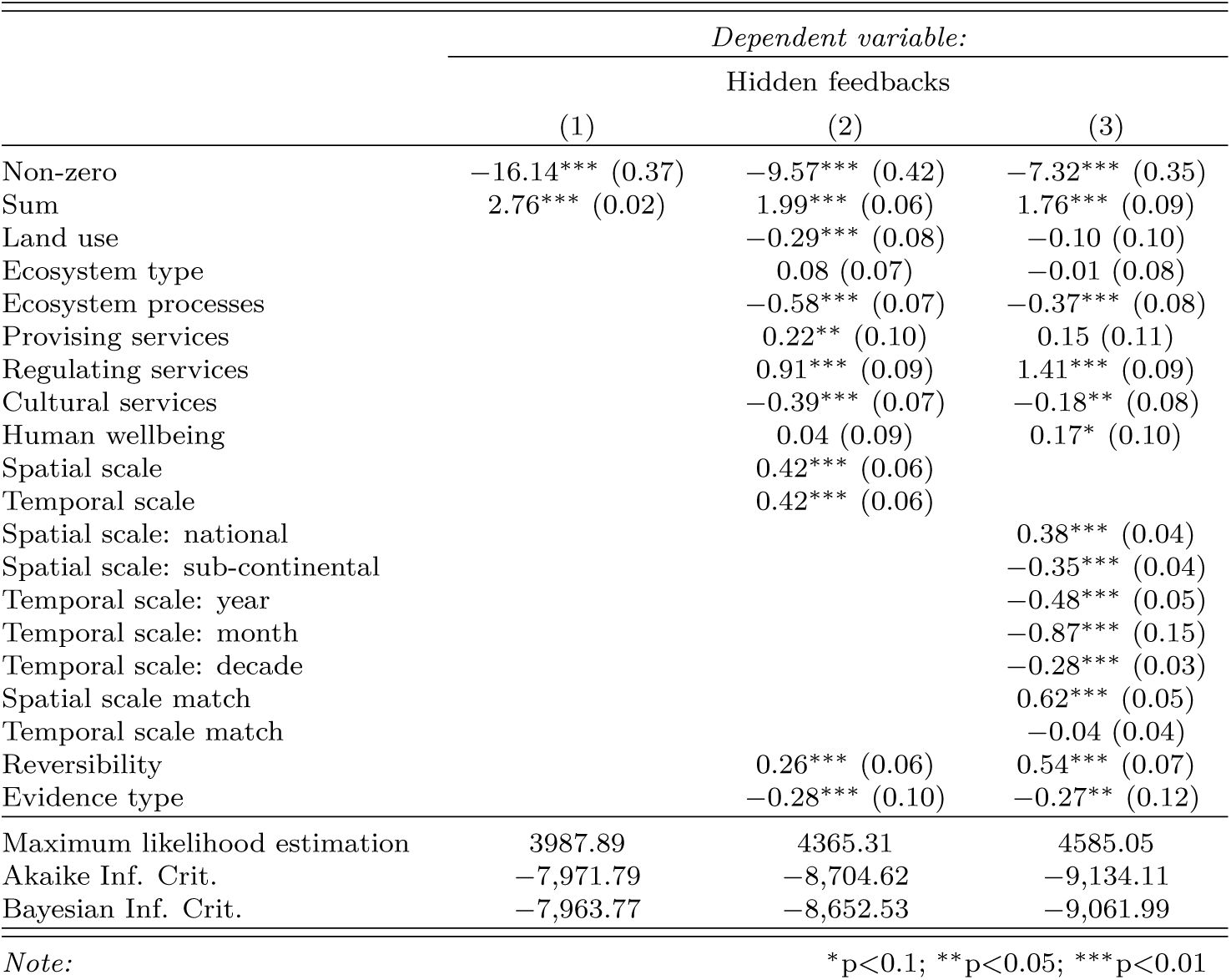
Exponential random graph models for *hidden feedbacks*

|  | *Dependent variable:* |  |  |
| --- | --- | --- | --- |
|  | Hidden feedbacks |  |  |
|  | (1) | (2) | (3) |
| Non-zero | −16.14*** (0.37) | −9.57*** (0.42) | −7.32*** (0.35) |
| Sum | 2.76*** (0.02) | 1.99*** (0.06) | 1.76*** (0.09) |
| Land use |  | −0.29*** (0.08) | −0.10 (0.10) |
| Ecosystem type |  | 0.08 (0.07) | −0.01 (0.08) |
| Ecosystem processes |  | −0.58*** (0.07) | −0.37*** (0.08) |
| Provising services |  | 0.22** (0.10) | 0.15 (0.11) |
| Regulating services |  | 0.91*** (0.09) | 1.41*** (0.09) |
| Cultural services |  | −0.39*** (0.07) | −0.18** (0.08) |
| Human wellbeing |  | 0.04 (0.09) | 0.17* (0.10) |
| Spatial scale |  | 0.42*** (0.06) |  |
| Temporal scale |  | 0.42*** (0.06) |  |
| Spatial scale: national |  |  | 0.38*** (0.04) |
| Spatial scale: sub-continental |  |  | −0.35*** (0.04) |
| Temporal scale: year |  |  | −0.48*** (0.05) |
| Temporal scale: month |  |  | −0.87*** (0.15) |
| Temporal scale: decade |  |  | −0.28*** (0.03) |
| Spatial scale match |  |  | 0.62*** (0.05) |
| Temporal scale match |  |  | −0.04 (0.04) |
| Reversibility |  | 0.26*** (0.06) | 0.54*** (0.07) |
| Evidence type |  | −0.28*** (0.10) | −0.27** (0.12) |
| Maximum likelihood estimation | 3987.89 | 4365.31 | 4585.05 |
| Akaike Inf. Crit. | −7,971.79 | −8,704.62 | −9,134.11 |
| Bayesian Inf. Crit. | −7,963.77 | −8,652.53 | −9,061.99 |

*Note:*
\*p<0.1; \*\*p<0.05; \*\*\*p<0.01

